# Liquid condensation drives telomere clustering during ALT

**DOI:** 10.1101/633040

**Authors:** Huaiying Zhang, Michel Liu, Robert Dilley, David M. Chenoweth, Roger A. Greenberg, Michael A. Lampson

## Abstract

Telomerase-free cancer cells employ a recombination-based alternative lengthening of telomeres (ALT) pathway that depends on <u>A</u>LT-associated <u>p</u>romyelocytic leukemia (PML) nuclear <u>b</u>odies (APBs), whose function is unclear. We find that APBs behave as liquid condensates, suggesting two potential mechanisms to promote telomere elongation: condensation to enrich DNA repair factors for telomere synthesis and coalescence to cluster telomeres to provide repair templates. Using chemically-induced dimerization, we show that telomere sumoylation nucleates APB condensation via SUMO-SIM (SUMO interaction motif) interactions and clusters telomeres. The induced APBs lack DNA repair factors, indicating that these factors are clients recruited to the APB scaffold rather than components that drive condensation. Telomere clustering, however, relies only on liquid properties of the condensate, as an alternative condensation chemistry also induces clustering. Our results demonstrate how the material properties and chemical composition of APBs independently contribute to ALT, suggesting a general framework for how liquid condensates promote cellular functions.

## Introduction

Telomeres are repetitive sequences at chromosome ends that shorten with each division in cells that lack a telomere maintenance mechanism. Critical telomere shortening induces replicative senescence or apoptosis (Harley et al., 1990), whereas cancer cells maintain proliferation potential by actively elongating their telomeres. The majority of human cancer cells re-activate the enzyme telomerase, but a significant fraction (10-15%) employ an alternative lengthening of telomeres (ALT) pathway that involves DNA recombination and repair to maintain telomere length (Dilley and Greenberg, 2015; Lazzerini-Denchi and Sfeir, 2016; Sobinoff and Pickett, 2017). The molecular mechanisms underlying ALT are unclear, but one unique characteristic is the presence of APBs, a class of <u>A</u>LT telomere-associated <u>p</u>romyelocytic leukemia (PML) nuclear <u>b</u>odies used for ALT diagnosis (Yeager et al., 1999). PML nuclear bodies are dynamic structures in the nucleus that transiently sequester up to 100 different proteins that are implicated in many cellular functions including tumor suppression, DNA replication, gene transcription, DNA repair, viral pathogenicity, cellular senescence and apoptosis (Lallemand-Breitenbach and de The, 2010). While APBs are proposed to be sites of telomere recombination during ALT, the precise functions of these specialized PML nuclear bodies are poorly understood. Inhibiting APB formation by knocking down PML protein, an essential component of PML nuclear bodies, leads to telomere shortening (Draskovic et al., 2009; Osterwald et al., 2015), indicating that APBs are essential for ALT telomere maintenance. In addition to typical PML nuclear body components, APBs contain proteins involved in homologous recombination such as replication protein A (RPA), Rad51, and breast cancer susceptibility protein 1 (BRCA1) (Nabetani and Ishikawa, 2011), which suggests that APBs promote telomere synthesis. Indeed, new telomere DNA synthesis has been detected in APBs (Cho et al., 2014; Chung et al., 2011; O’sullivan et al., 2014; Sahin et al., 2014; Zhang et al., 2019). Telomeres also cluster within APBs, as another unique feature of ALT, presumably to provide repair templates for telomere DNA synthesis. Many functionally distinct proteins can initiate APB assembly, leading to the proposal of a multiple-pathway model (Chung et al., 2011), as suggested by an RNA interference screen that identified close to thirty proteins that affect APB formation, including proteins involved in telomere and chromatin organization, protein sumoylation, and DNA repair (Osterwald et al., 2015). Given such complexity, the mechanisms governing APB assembly and function remain unclear, and limitations include lack of a conceptual model for how they form and lack of tools to manipulate the process for cell biological analyses. We previously showed that introducing DNA damage on telomeres leads to APB formation, telomere clustering within the induced APBs, and telomere elongation (Cho et al., 2014). While DNA damage from either replication stress or DSBs at telomeres can trigger APB formation (O’Sullivan NSMB 2014; Cho et al. Cell 2014), the physical mechanisms underlying such clustering are unknown.

Many nuclear bodies and membrane-free organelles – such as P granules, nucleoli, signaling complexes, and stress granules (Altmeyer et al., 2015; Brangwynne et al., 2011, 2009; Patel et al., 2015a; Su et al., 2016) – assemble by liquid-liquid phase separation, in which proteins and/or nucleic acids separate from the surrounding milieu and form a condensed liquid phase (Banani et al., 2017). Components of these condensates are highly concentrated but can dynamically exchange with the diluted phase. Liquid phase separation provides a mechanism for organizing matter in cells, particularly protein interaction networks that do not form stable complexes with fixed stoichiometry. Notably, such stable complexes are relatively rare, and protein-protein interactions are dominated by weak interactions (Hein et al., 2015). In vitro reconstitution has provided valuable insights on how those weak interactions drive the condensation process, but little is known about how liquid phase separation, particularly the unique liquid properties of the resulting condensates, promote cellular functions.

Using a chemical inducer of protein dimerization, we induce de novo APB formation in live cells and provide evidence that APBs assemble via liquid-liquid phase separation, driven by SUMO-SIM interactions. We find that the coalescence of APB liquid droplets drives telomere clustering, depending only on the liquid properties of APBs. Overall, this work provides tools to manipulate APB assembly, a conceptual model for APB assembly via liquid-liquid phase separation, and insight into how APB condensation contributes to ALT.

## Results

### SUMO-SIM interactions drive APB liquid condensation to cluster telomeres

Previously, we introduced DNA damage on telomeres in ALT cells by fusing the FokI nuclease to the telomere binding protein TRF1, which induced APB formation, telomere clustering within APBs, and telomere elongation (Cho et al., 2014). With this assay, we observed that APBs exhibit liquid behavior, including coalescence after colliding (Figure 1A, B) and dynamic exchange of components within APBs and with the surrounding nucleoplasm, as shown by fluorescence recovery after photobleaching (Figure 1C). These phenomena are characteristics of liquid condensates formed by liquid-liquid phase separation, leading us to hypothesize that APBs are liquid droplets condensed on telomeres after DNA damage as a mechanism for telomere clustering and elongation. The liquid nature of APBs would promote telomere clustering via coalescence, and the condensates may serve as platforms to concentrate DNA repair factors to aid telomere synthesis. The switch-like self-assembly and disassembly of liquid droplets would allow APBs to rapidly nucleate as telomeres shorten and subsequently dissolve by reversing the nucleation signal.

**Figure 1.**
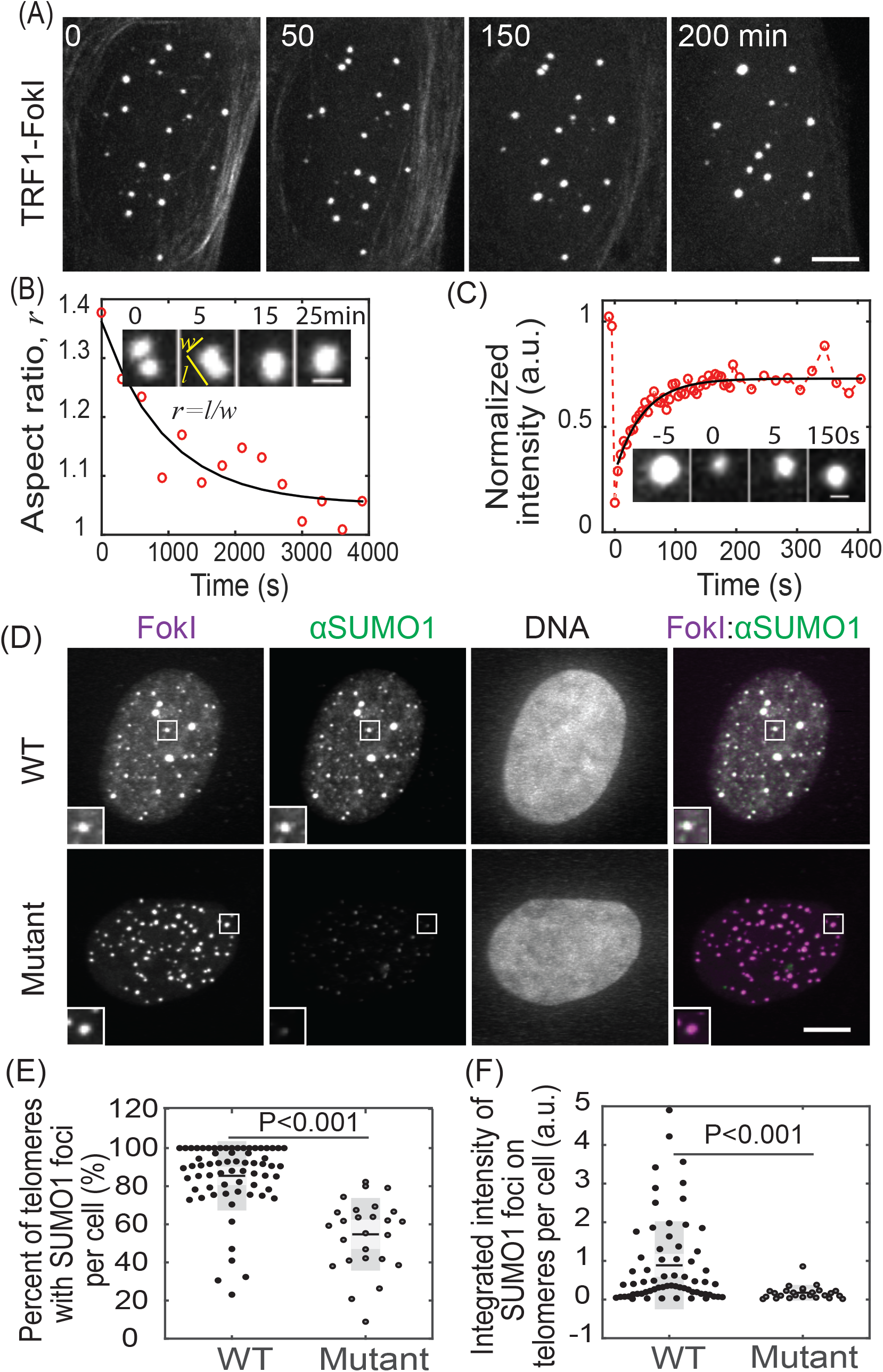
APBs exhibit liquid behavior and concentrate SUMO. APB formation was induced by creating DNA damage on telomeres with TRF1-FokI. **(A-B)** Cells were imaged live starting 1 hour after triggering mCherry-TRF1-FokI import into the nucleus. Images show clustering of TRF1 foci (A) and fusion (B, insets), quantified by change in aspect ratio (defined as length/width) over time (exponential fit: 15 min half time). **(C)** Fluorescence recovery after photobleaching (FRAP) of DNA damage-induced APBs. Insets shows a single APB, intensity normalized to the first time point, exponential fit: 30 s recovery half time. **(D-F)** SUMO1 immunofluorescence for cells expressing TRF1-FokI or a nuclease-dead mutant. The overlay of FokI (purple) and SUMO1 (green) appears white (D, insets two times enlarged). Graphs show the percent of telomeres with SUMO1 foci and the integrated intensity of SUMO1 foci on telomeres. Each data point represents one cell from two biological replicates, black lines mean, gray bars 95% confidence interval. Scale bars 5 μm (A, D) or 1 μm (B, C). Also see Figure 1-figure supplement 1.

We considered the possibility that sumoylation of telomere-binding proteins (e.g., shelterin complex) triggers APB condensation, driven by multivalent SUMO-SIM interactions. Many APB components are SUMOylated, contain SIM domains, or both (Chung et al., 2011; Potts and Yu, 2007; Shen et al., 2006; Shima et al., 2013), and sumoylation of telomere proteins is required for APB formation (Potts and Yu, 2007). Furthermore, synthetic SUMO and SIM peptides can drive liquid droplet formation in vitro (Banani et al., 2016a). These findings are consistent with a model in which SUMO-SIM interactions on telomere binding proteins cooperate during phase separation to drive telomere coalescence into ABPs. DNA damage responses triggered by telomere shortening would be a stimulus to induce SUMOylation. Conversely, desumoylation after telomere elongation would lead to APB dissolution. Supporting this idea, we observed enrichment of both SUMO1 and SUMO2/3 after DNA damage induced with FokI, but not with a FokI mutant that lacks nuclease activity (Figure 1D-F, Figure 1-figure supplement 1).

To test the hypothesis that telomere sumoylation drives APB condensation via SUMO-SIM interactions, we developed a protein dimerization approach to induce de novo APB formation on telomeres without DNA damage. To mimic sumoylation on telomeres and avoid overexpressing SUMO, we recruited SIM to telomeres with a chemical inducer of dimerization. We predicted that SIM recruited to telomeres can bring endogenous SUMO to telomeres to induce APB condensation. The chemical dimerizer consists of two linked ligands: trimethoprim (TMP) and Haloligand, and can dimerize proteins fused to the cognate receptors: *Escherichia coli* dihydrofolate reductase (eDHFR) and a bacterial alkyldehalogenase enzyme (Haloenzyme), respectively (Figure 2A). An advantage of this system is that it is reversible by adding excess TMP to compete for eDHFR, unlike other chemically induced dimerization systems such as rapamycin (Ballister et al., 2014; DeRose et al., 2013). We fused Haloenzyme to the telomere binding protein TRF1 to anchor it to telomeres and to GFP for visualization. SIM was fused to eDHFR and to mCherry. After adding the dimerizer to cells expressing Halo-GFP-TRF1 and SIM-mCherry-eDHFR, SIM was recruited to telomeres, which resulted in enrichment of both SUMO1 and SUMO2/3 on telomeres (Figure 2B-D, Figure 2-figure supplement 1). To confirm that enrichment of SUMO is indeed based on SUMO-SIM interaction, we used a SIM mutant that cannot interact with SUMO (Banani et al., 2016b). As predicted, the SIM mutant was recruited to telomeres without SUMO enrichment. To confirm that the sites of SIM recruitment are telomeres, we used fluorescence in situ hybridization (FISH) to visualize telomere DNA directly and observed colocalization of SIM with telomere signal (Figure 2E).

**Figure 2.**
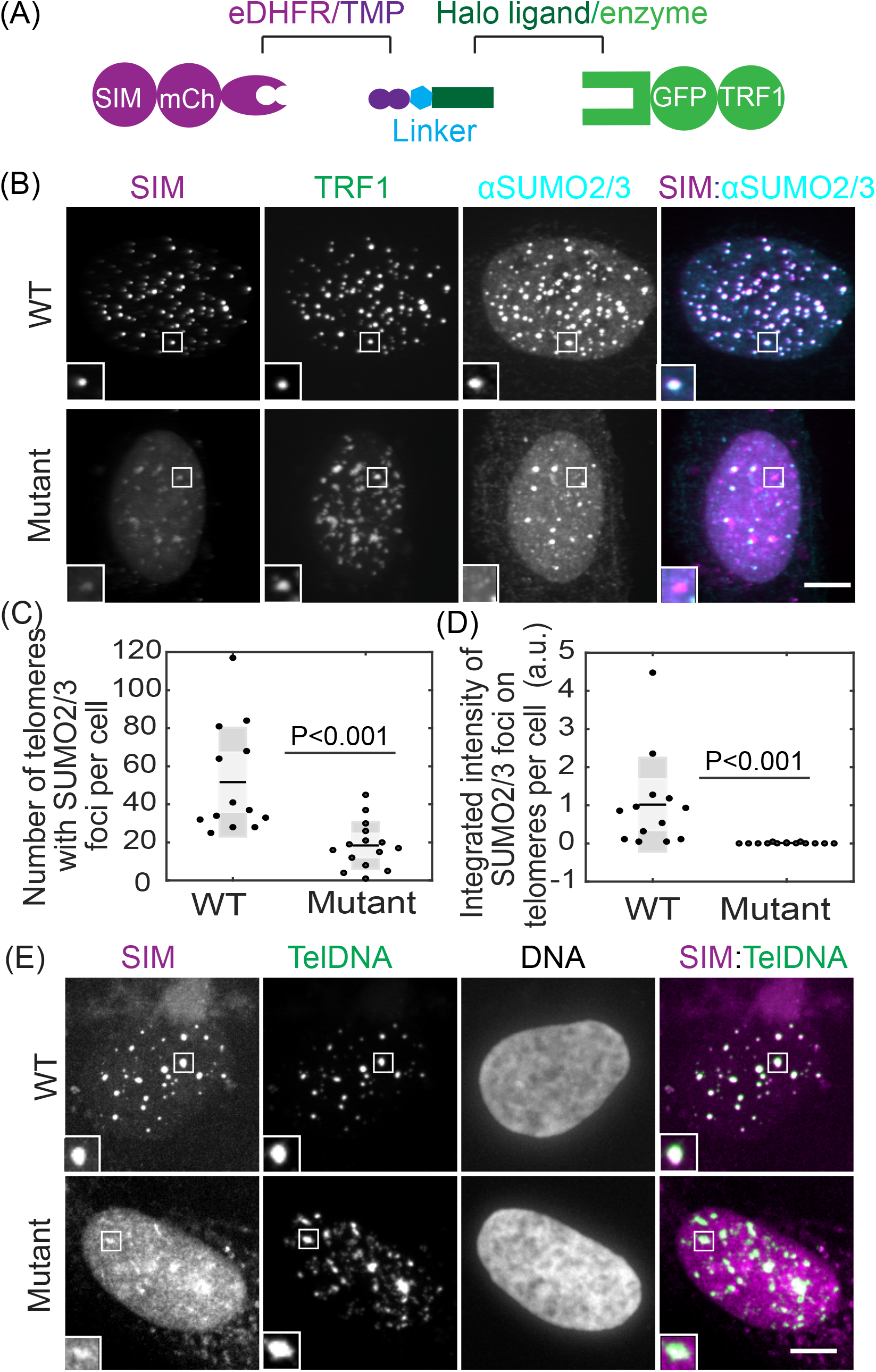
Recruiting SUMO to telomeres through SIM with a chemical dimerizer. **(A)** Dimerization schematic: SIM is fused to mCherry and eDHFR, and TRF1 is fused to Halo and GFP. The dimerizer is TNH: TMP(trimethoprim)-NVOC (6-nitroveratryl oxycarbonyl)-Halo (Zhang et al., 2017). **(B-D)** Cells expressing SIM-mCherry-DHFR (WT) or a SIM mutant that cannot interact with SUMO, together with Halo-GFP-TRF1, were incubated with TNH before fixing and staining for SUMO2/3. The overlay of SIM (purple) and SUMO2/3 (cyan) appears white (B, insets two times enlarged). Graphs show the number of telomeres with SUMO2/3 foci and the integrated intensity of SUMO2/3 foci on telomeres. Each data represents one cell from two biological replicates, black lines mean, gray bars 95% confidence interval. **(E)** Telomere FISH images after recruiting SIM or SIM mutant to telomeres. The overlay of SIM (purple) and telomere DNA FISH (green) appears white. Scale bars 5 μm. Also see Figure 2-figure supplement 1.

To directly test whether SIM recruitment leads to liquid condensation on telomeres, we used live imaging to monitor TRF1 and SIM signals over time (Movie 1). We observed that after SIM recruitment, both SIM and TRF1 foci became brighter and bigger (Figure 3A, B), as predicted for liquid droplet nucleation and growth. In addition, both SIM and TRF1 foci rounded up, indicating formation of liquid condensates. Such liquid behavior is also shown by fusion events and the dynamic exchange of components, similar to DNA damage-induced foci (Figure 3D, E). Droplet fusion also drove telomere clustering, leading to reduced telomere number over time (Figure 3C), although as in previous studies, we cannot differentiate telomeres from extrachromosomal telomere DNA that exists in ALT cells (Cho et al., 2014). In contrast, a SIM mutant that cannot interact with SUMO was recruited to telomeres after dimerization, but did not induce condensation or telomere clustering (Movie 2, Figure 3-figure supplement 1). Overall, these findings support our hypothesis that condensation is driven by SUMO-SIM interactions.

**Figure 3.**
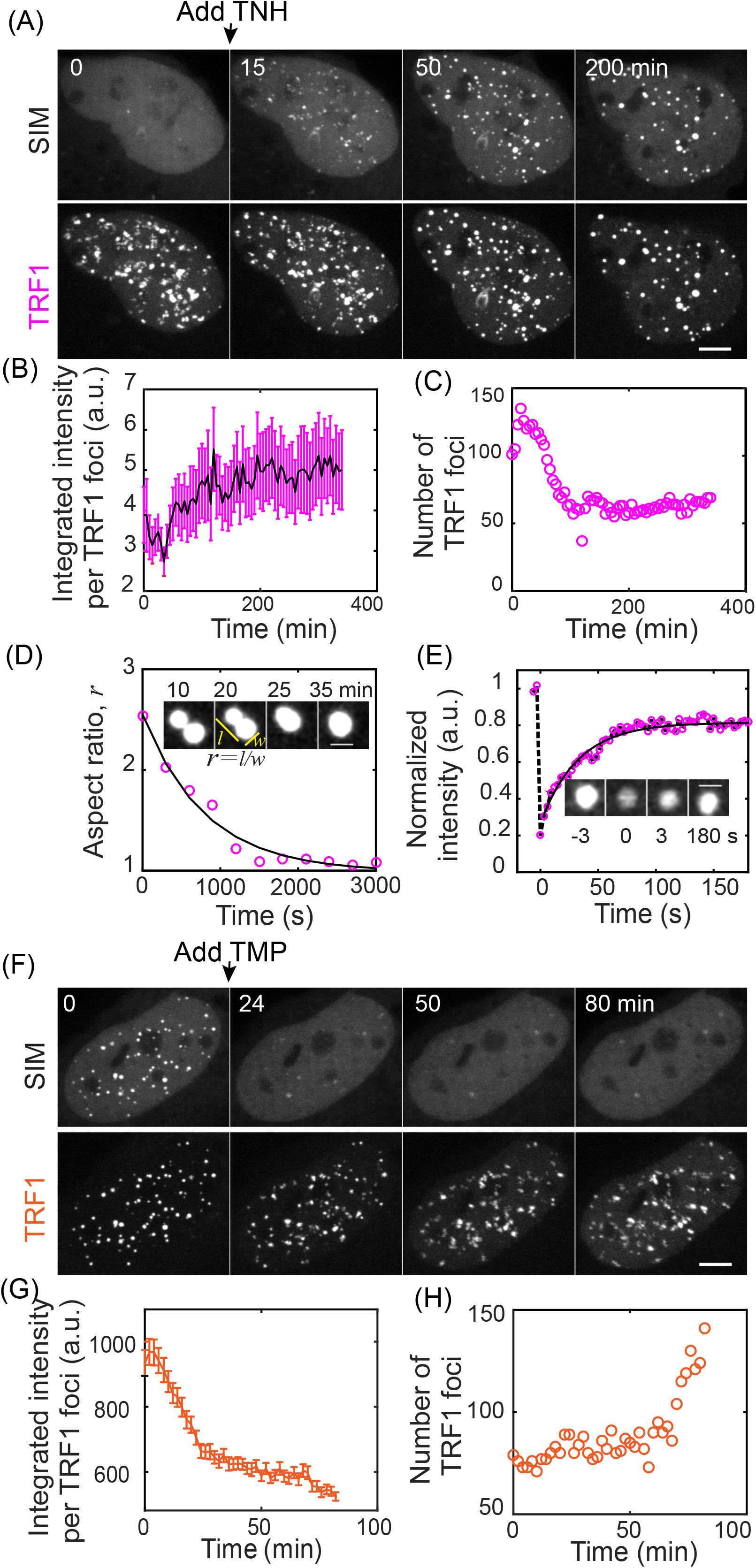
SUMO-SIM interactions drive liquid condensation and telomere clustering. **(A-D)** TNH was added to cells expressing SIM-mCherry-DHFR and Halo-GFP-TRF1 after the first time point to induce dimerization. Graphs show mean integrated intensity per TRF1 foci (B, error bars SEM) and number of TRF1 foci (C) over time. Insets (D) show an example of a fusion event, with the change in aspect ratio quantified (exponential fit, decay time 13 min). **(E)** FRAP of dimerization-induced condensates. Intensity is normalized to the first time point, exponential fit: 22 s recovery half time. **(F-H)** After dimerization induced by TNH in cells expressing SIM-mCherry-DHFR and Halo-GFP-TRF1, TMP was added to release SIM from telomeres. Scale bars 5 μm. Also see Figure 3-figure supplement 1.

Our phase transition model predicts that reversal of the nucleation signal will result in the dissolution of condensates. To test this prediction, we first formed condensates on telomeres by SIM recruitment and then added free TMP to compete with the dimerizer for eDHFR binding to reverse dimerization (Ballister et al., 2014). Condensation and telomere clustering were reversed as the intensity decreased in the foci while increasing in the nucleoplasm (Movie 3, Figure 3F, G), and telomere number increased (Figure 3H), consistent with our model.

### Functional contributions of APB condensates

APB condensates could promote homology-directed telomere DNA synthesis in ALT by either or both of two mechanisms: 1) concentrating DNA repair factors on telomeres through APB condensation, 2) clustering telomeres for repair templates through APB coalescence. The first mechanism is an example of compositional control of phase-separated condensates, which can be described with a scaffold-client model (Banani et al., 2016a). Scaffold components interact with each other to drive condensation, and clients are recruited to the condensate. Functionally, scaffold components provide a platform for concentrating clients together for a cellular function. Clients can be recruited via direct interactions with scaffold components or via additional signaling such as post-translational modifications.

To determine whether APB condensates follow a scaffold-client model, we examined DNA repair factors and PML protein, whose localization on telomeres defines APBs. Recruiting SIM to telomeres increased colocalization of PML with telomeres, compared to control cells where SIM was not recruited (Figure 4A-C). Together with our previous findings that the dimerization-induced condensates contain other known components of APBs – SUMO (Figure 2B-D, Figure 2-figure supplement 1), telomere DNA (Figure 2E) and TRF1 (Figure 3) – this result indicates that the induced condensates are indeed APBs with PML as a scaffold component. Such an increase in PML localization to telomeres was not seen when the SIM mutant was recruited, agreeing with the hypothesis that SUMO-SIM interactions drive APB condensation. As potential clients, we looked at proteins involved in the DNA damage response and repair pathways: 53BP1, PCNA, and POLD3, which localize to APBs induced by DNA damage (Cho et al., 2014; Dilley et al., 2016). None of these factors was recruited after dimerization-induced condensation (Figure 4D-F, Figure 4-figure supplement 1), indicating that they are client molecules recruited to the APB scaffold, independent of condensation, via additional signaling.

**Figure 4.**
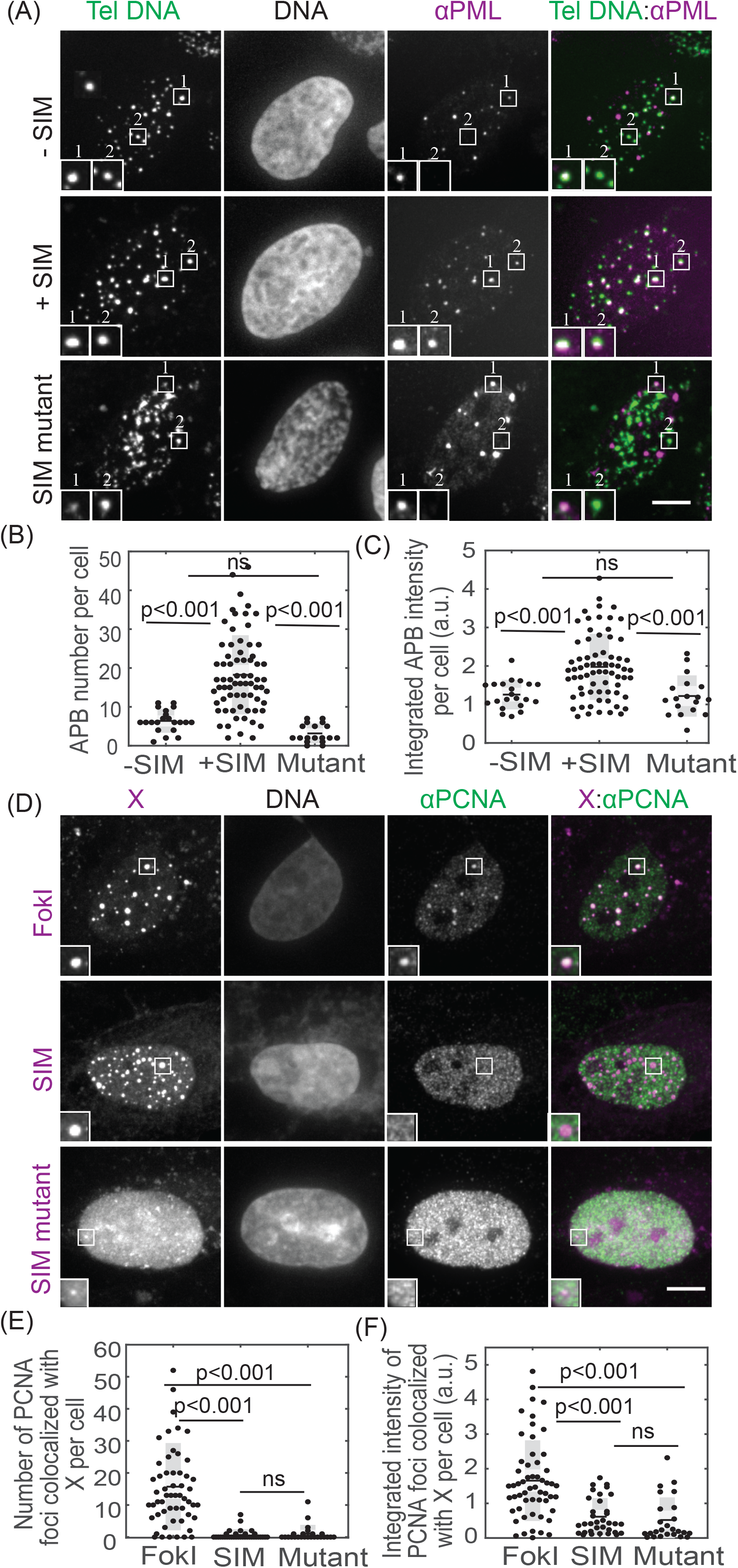
Condensates contain APB scaffold components but not DNA repair factors. **(A-C)** FISH of telomere DNA and immunofluorescence of PML for cells with or without SIM recruited to telomeres or with SIM mutant recruited to telomeres. The overlay of PML (purple) and telomere DNA (green) appears white (A, insets two times enlarged), indicating APBs with PML nuclear bodies localized to telomeres. Graphs show APB number and integrated APB intensity per cell. **(D-F)** Immunofluorescence of PCNA for cells with FokI-induced damage or with SIM or SIM mutant recruited. In representative images (D, insets two times enlarged), X indicates FokI, SIM or SIM mutant, and colocalization with PCNA appears white in overlay images (right panels). Graphs show number of PCNA foci colocalized with FokI, SIM, or SIM mutant and integrated intensity. Each data point (B, C, E, F) represents one cell from two biological replicates, black line mean, gray bar 95% confidence interval. Scale bars 5 μm. Also see Figure 4-figure supplement 1.

Our model predicts that the ability to cluster telomeres relies on the liquid material properties of APBs and not on specific scaffold or client proteins. To test this prediction, we aimed to induce non-APB liquid droplets with a different chemistry on telomeres and determine whether they can cluster telomeres. Besides multivalent interactions between modular interaction pairs such as SUMO and SIM, another way of driving condensation is through interactions between disordered or no complexity protein domains that behave like flexible polymers (Elbaum-Garfinkle et al., 2015a; Lin et al., 2015; Nott et al., 2015; Patel et al., 2015b; Zhang et al., 2015). We selected the arginine/glycine-rich (RGG) domain from the P granule component LAF-1, which forms liquid condensates in vitro and in vivo (Elbaum-Garfinkle et al., 2015b; Schuster et al., 2018). Recruiting RGG to telomeres resulted in condensation as shown by the increase in telomere foci intensity (Movie 4, Figure 5A, B). The induced condensates exhibited liquid behavior such as the ability to fuse, which led to telomere clustering as shown by the decrease in telomere foci over time (Figure 5C, D). We also confirmed that the RGG condensates were indeed on telomeres, and did not increase PML protein on telomeres compared with cells without RGG recruited (Figure 5 E-G), indicating the induced condensates are not APBs. These results support our model that liquid condensation drives telomere clustering independent of specific protein components of the condensates.

**Figure 5.**
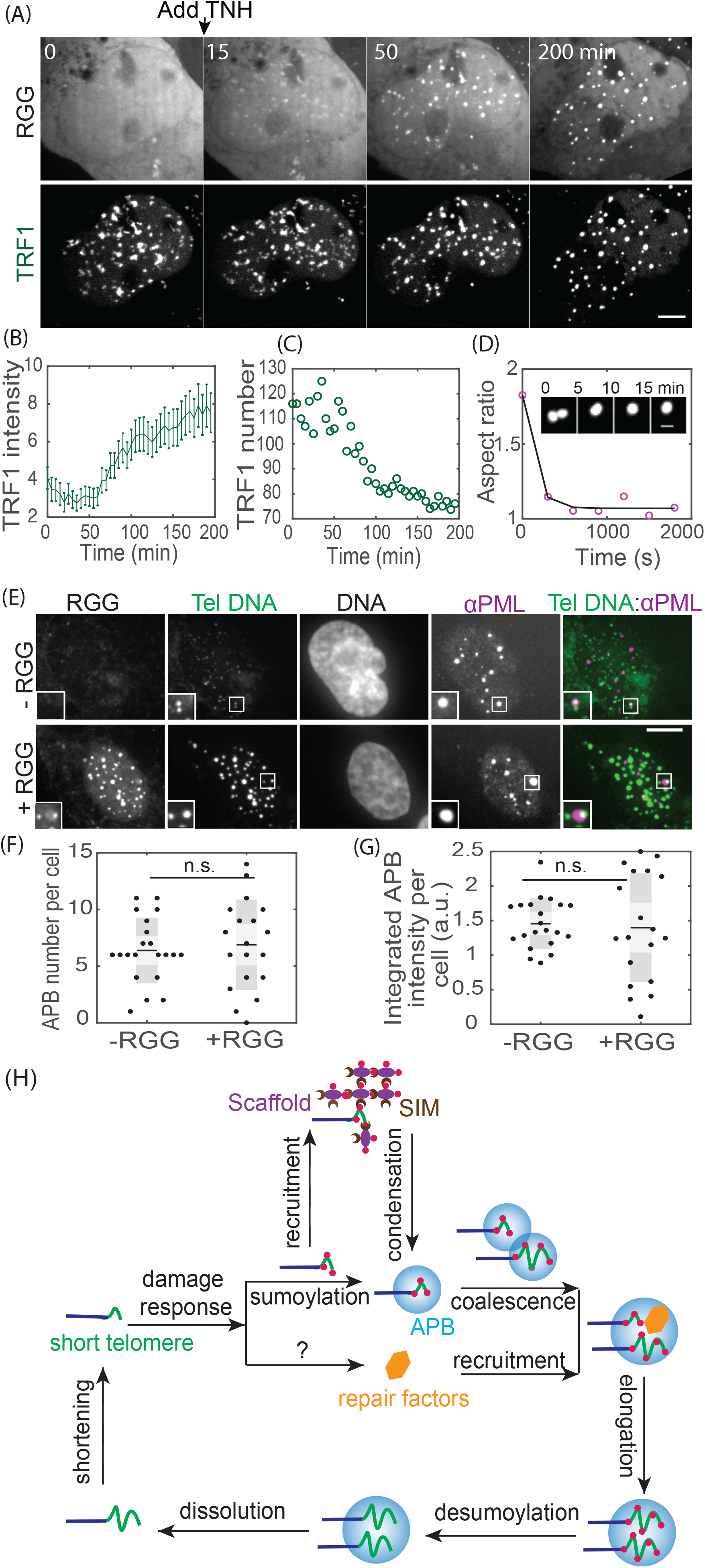
Non-APB condensation on telomeres drives telomere clustering. **(A-D)** TNH was added to cells expressing RGG-mCherry-RGG-eDHFR and Halo-GFP-TRF1 to induce dimerization and condensation. Graphs show integrated intensity per TRF1 foci (B, error bars SEM) and number of TRF1 foci (C) over time. Insets (D) show an example of a fusion event, with the change in aspect ratio quantified (exponential fit, decay time 6 min). **(E-G)** FISH of telomere DNA and immunofluorescence of PML for cells with or without RGG recruitment. In representative images (E) the overlay of PML (purple) and telomere DNA (green) appears white, indicating APBs with PML nuclear bodies localized to telomeres. Insets (two times enlarged) show two telomere foci, one with an APB and one without, indicating the basal level of APBs in these cells. Graphs show APB number per cell and integrated APB intensity per cell. Each data point (F, G) represents one cell from two biological replicates, black line mean, gray bar 95% confidence interval. **(H**) Model for APB condensation and function. Telomere shortening (or replication stress) triggers a DNA damage response, where telomere sumoylation nucleates APB condensation and drives telomere clustering while another aspect of the damage response pathway recruits DNA repair factors to APB condensates. Together the clustered telomeres and enriched DNA repair factors within APBs lead to homology-directed telomere synthesis in ALT. Scale bars 5 μm.

## Discussion

We propose a liquid-liquid phase separation model for APB assembly triggered by telomere sumoylation via SUMO-SIM interactions as part of a DNA damage response at telomeres(Figure 5H). We induced APB condensation by recruiting SIM to telomeres using a chemical dimerizer, independent of DNA damage. Conversely, we find that releasing SIM from telomeres reverses APB condensation. These findings indicate that APB condensates are nucleated on telomeres via sumoylation and dissolved via desumoylation. Sumoylation has long been observed as part of the DNA damage response (Hendriks and Vertegaal, 2015), and PML nuclear bodies are also implicated in DNA repair (Dellaire and Bazett-Jones, 2004), though the molecular mechanisms of both remain unclear. Our observation that sumoylation nucleates APB condensates as a mechanism for telomere elongation may lead to future insights on the roles of sumoylation and PML bodies in DNA repair in other contexts (Sarangi and Zhao, 2015; Xu et al., 2003).

The induced condensates contain the APB signature component PML but not DNA repair factors such as 53BP1, PCNA and POLD3 (Figure 4, Figure 4-figure supplement 1), indicating that the repair factors are client molecules recruited to the APB scaffold by DNA damage response signaling other than the telomere sumoylation that nucleates APBs (Figure 5H). Other aspects of sumoylation, such as protein conformational changes, are not mimicked by our SIM recruitment approach and may be important for recruiting DNA repair factors. In addition to telomere binding proteins, many DNA repair factors are also sumoylated including 53BP1 and PCNA (Garvin and Morris, 2017). Sumoylation of those DNA repair factors, not captured in our dimerization approach, may be required for recruitment to APBs. It is also possible that other posttranslational modifications (PTMs) are required to regulate interactions of DNA repair factors with APB components. Indeed, the DNA damage response is a complex process that involves a multitude of PTMs including phosphorylation, ubiquitylation, sumoylation, neddylation, poly (ADP-ribosyl)ation, acetylation, and methylation of chromatin and chromatin-associated proteins (Dantuma and van Attikum, 2016). It is still unclear how those PTMs are spatially and temporally regulated to work together in DNA repair. Here we show that sumoylation is responsible for nucleating APBs, and future studies revealing what signaling is required for recruitment of client molecules to the APB scaffold will provide insights on how sumoylation together with other PTMs promotes telomere DNA synthesis in ALT and in DNA repair more broadly.

We showed that coalescence of APB liquid droplets drives telomere clustering (Figure 3A-E), which is thought to provide repair templates for homology-directed telomere DNA synthesis in ALT. ALT cells contain extrachromosomal telomere DNAs (ECTRs) that may either be linear or circular, but their functional contribution to ALT is unknown (Cesare and Griffith, 2004). They share sequence identity with telomeres and cannot be differentiated with our TRF1 probe or other labeling techniques targeting telomere DNA sequence. Therefore, the clustering we observe may involve APBs nucleated on both telomeres and ECTRs. Since ECTRs are more mobile, they may be more efficient in clustering with telomeres to provide repair templates. Further studies dissecting the role of ECTRs in telomere clustering would increase our understanding of templating in ALT. We also demonstrated that the ability to cluster telomeres depends only on the liquid properties of APB condensates, not their chemical composition (Figure 5). This finding provides an opportunity to target the physical-chemical properties of APBs for cancer therapy in ALT without affecting function of APB components in normal cells.

Liquid-liquid phase separation can contribute to cellular functions by multiple mechanisms. For example, the high sensitivity of the phase separation process to environmental factors makes it ideal for sensing stress (Munder et al., 2016; Riback et al., 2017), and concentrating and confining molecules into one compartment can increase the kinetics of biochemistry (Case et al., 2019). The hallmark of such phase separation is the liquid properties of the resulting condensates, which have been carefully characterized in reconstituted systems. The functional significance of these in vitro findings in cells have been widely implied but not demonstrated yet (Shin and Brangwynne, 2017). With dimerization-induced condensation, we show that the liquid properties of APB condensates drive telomere clustering, independent of condensate chemistry. Our findings may represent a general strategy for reversible genome organization, such as clustering of gene loci for transcription and DNA repair, and suggest a dual function model for chromatin condensates: concentrating factors for biochemistry through composition control while clustering distinct chromatin domains via coalescence.

## Acknowledgments

We thank Stephanie Weber for sharing MATLAB code for analyzing foci intensity in 3D and Michael Rosen and Benjamin Schuster for sharing plasmids. We thank members of the Greenberg lab and Lampson lab for helpful discussions. This work was supported by the National Institutes of Health (GM122475 to M.A.L., GM118510 to D.M.C., U54-CA193417 to Physical Sciences Oncology Center at Penn, 1K22CA237632-01 to H.Z., GM101149 and CA17494 to R.A.G.).

---

**Figure 1-figure supplement 1.**
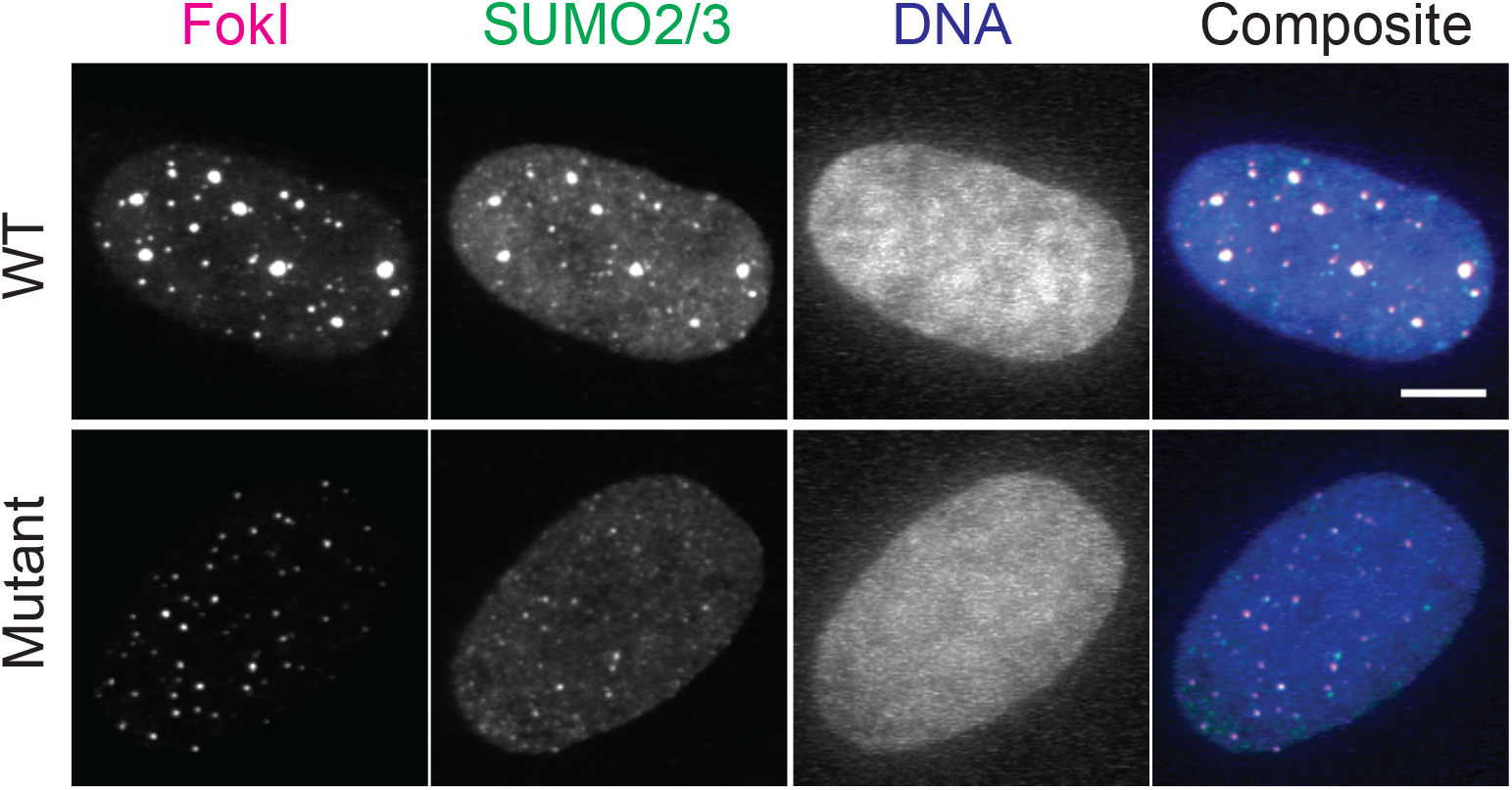
SUMO2/3 is enriched on telomeres after DNA damage. Immunofluorescence images of SUMO2/3 for cells with FokI or a nuclease dead FokI mutant targeted to telomeres. Scale bars 5 μm.

**Figure 2-figure supplement 1.**
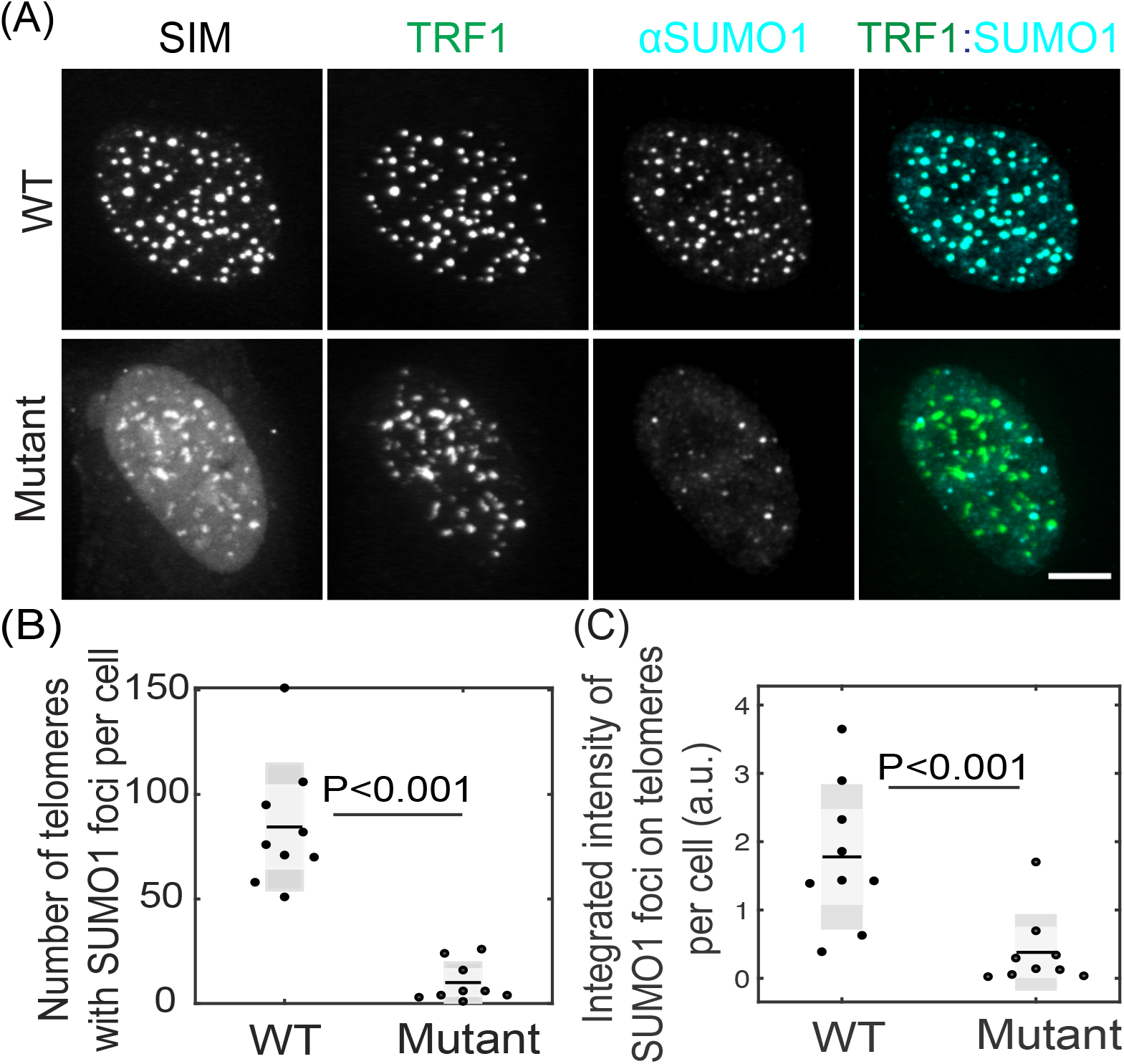
SUMO1 is enriched on telomeres after SIM recruitment. Immunofluorescence images of SUMO1 after recruiting SIM or SIM mutant to telomeres (A), and quantification of the number of telomeres with SUMO1 foci per cell (B) and the integrated intensity of SUMO1 foci on telomeres per cell (C). Each data point in (B) and (C) represents one cell from two biological replicates, black line represents the average value, and gray bar represents 95% confidence interval. Scale bars 5 μm.

**Figure 3-figure supplement 1.**
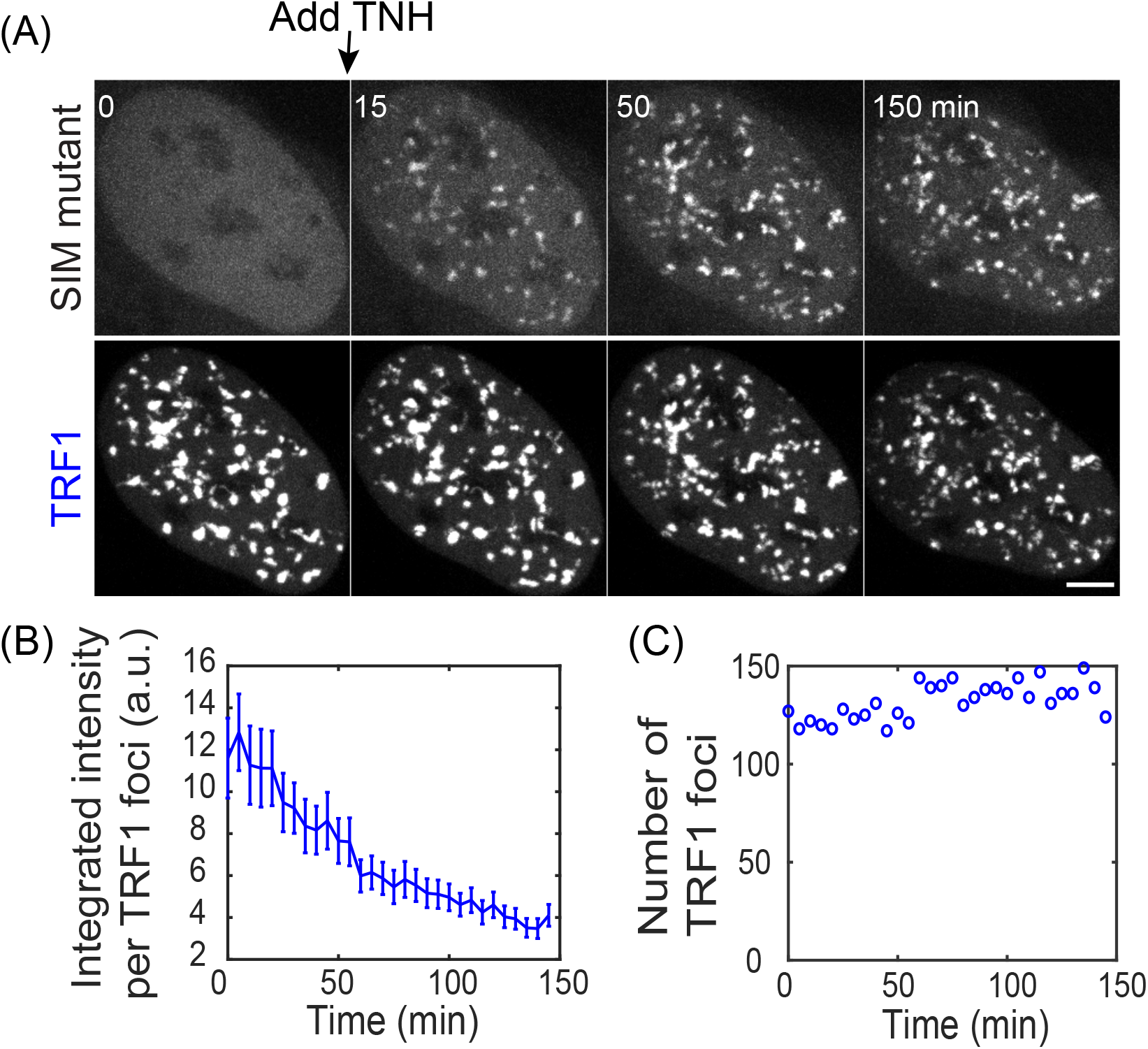
SIM mutant recruited to telomeres cannot induce condensation and clustering. TNH was added to cells expressing SIM mutant-mCherry-DHFR and Halo-GFP-TRF1 after the first time point to induce dimerization. Graphs show mean integrated intensity per TRF1 foci (B, error bars SEM) and number of TRF1 foci (C) over time. In contrast to SIM recruitment, telomere number stayed unchanged and the intensity was not increased, but decreased due to photobleaching

**Figure 4-figure supplement 1.**
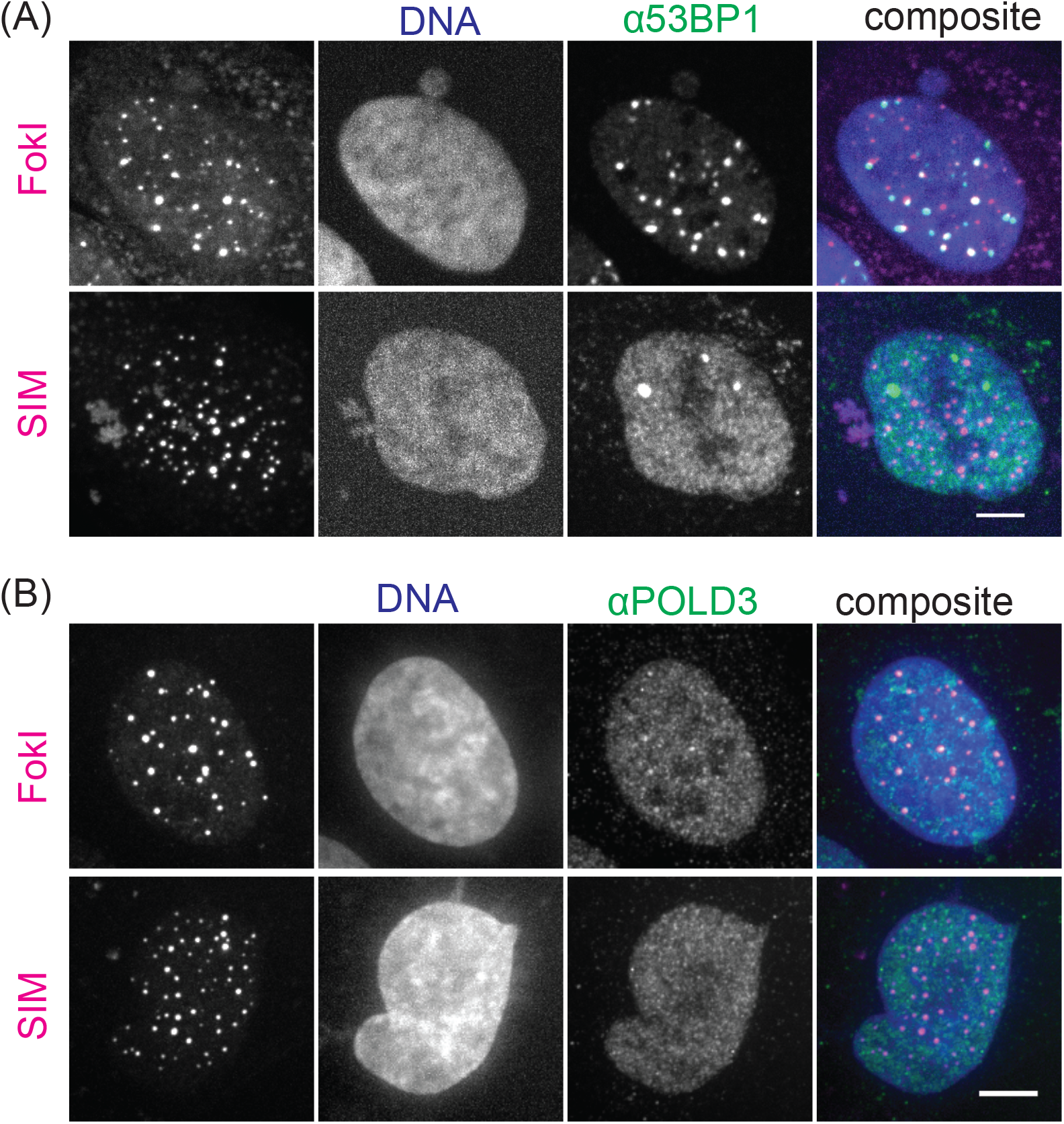
Unlike damaged-induced APBs, dimerization-induced condensates do not enrich 53BP1 or POLD3. **(A)** Immunofluorescence images of 53BP1 (A) or POLD3 (B) on telomeres with FokI tethered or SIM recruited. Scale bars 5 μm.

**Movie 1. Recruit SIM to telomeres.** Movie for Figure 3A. Left: SIM-mCherry-eDHFR, middle: Halo-GFP-TRF1, right: composite of SIM (magenta) and TRF1 (green). TNH was added to cells after the first time point to induce dimerization. Yellow box highlights a fusion event.

**Movie 2. Recruit SIM mutant to telomeres.** Movie for Figure 3-figure supplement 1. Left: SIM mutant-mCherry-eDHFR, middle: TRF1-GFP-Halo, right: composite of SIM mutant (magenta) and TRF1 (green). TNH was added to cells after the first time point to induce dimerization.

**Movie 3. Release SIM from telomeres.** Movie for Figure 3F. Left: SIM-mCherry-eDHFR, middle: Halo-GFP-TRF1, right: composite of SIM (magenta) and TRF1 (green). Cells were incubated with TNH to induce dimerization for 2 hours before imaging. TMP was added after the first time point to release SIM from telomeres.

**Movie 4. Recruit RGG to telomeres.** Movie for Figure 5A.Left: RGG-mCherry-RGG-eDHFR, middle: Halo-GFP-TRF1, right: composite of RGG (magenta) and TRF1 (green). TNH was added to cells after the first time point to induce dimerization. Yellow box highlights a fusion event.

## Materials and Methods

### Plasmids

The plasmids for inducing DNA damage at telomeres (mCherry-ER-DD-TRF1-FokI or Fok1 mutant) were previously published (Cho et al., 2014). For recruiting SIM to telomeres, TRF1 was substituted for SPC25 in the published 3xHalo-GFP-SPC25 plasmid (Zhang et al., 2017). SIM (or SIM mutant) for SIM-mCherry-eDHFR is from plasmids gifted by Michel Rosen (Banani et al., 2016b). The RGG insert for RGG-mCherry-RGG-eDHFR is from a plasmid gifted by Benjamin Schuster (Schuster et al., 2018). The vector containing mCherry-eDHFR is from our published plasmid Mad1-mCherry-eDHFR (Zhang et al., 2017). All other plasmids in this study are derived from a plasmid that contains a CAG promoter for constitutive expression, obtained from E. V. Makeyev (Khandelia et al., 2011).

### Cell culture

All experiments were performed with U2OS acceptor cells, originally obtained from E.V. Makayev, Nanyang Technological University, Singapore(Khandelia et al., 2011). Cells were cultured in growth medium (Dulbecco’s Modified Eagle’s medium with 10% FBS and 1% penicillin–streptomycin) at 37 °C in a humidified atmosphere with 5% CO_2_. The TRF1 constructs (3xHalo-GFP-TRF1, 3xHalo-TRF1, or mCherry-ER-DD-TRF1-FokI) and the eDHFR constructs (SIM, SIM mutant, or RGG) were transiently expressed by transfection with Lipofectamine 2000 (Invitrogen) 24 hours prior to imaging, following the manufacturer’s protocol.

### Dimerization and damage on telomeres

To recruit proteins to telomeres, cells transfected with 3xHalo-GFP-TRF1 or 3xHalo-TRF1 and one of the mCherry-eDHFR plasmids (SIM, SIM mutant, or RGG) were treated with the dimerizer TNH: TMP(trimethoprim)-NVOC (6-nitroveratryl oxycarbonyl)-Halo (Zhang et al., 2017). For live imaging, 100 nM TNH was added directly to cells on the microscope stage. For IF or FISH, 100 nM TNH was added to cells and incubated for 2 hours before fixing. To induce damage on telomere in cells transfected with mCherry-ER-DD-TRF1-FokI, Shield-1 (Cheminpharma LLC) and 4-hydroxytamoxifen (4-OHT) (Sigma-Aldrich) at 1μM were added for one hour to allow TRF1 to enter the nucleus prior to live imaging or two hours prior to fixing, as previously described (Cho et al., 2014).

### Immunofluorescence (IF) and fluorescence in situ hybridization (FISH)

Cells were fixed in 4% formaldehyde for 10 min at room temperature, followed by permeabilization in 0.5% Triton X-100 for 10 min. Cells were incubated with primary antibody at 4°C in a humidified chamber overnight and then with secondary antibody for one hour at room temperature before washing and mounting. Primary antibodies were anti-SUMO1 (Ab32058, Abcam,1:200 dilution), anti-SUMO2/3 (Asm23, Cytoskleton, 1:200 dilution), anti-PCNA (P10, Cell Signaling, 1:1000 dilution), anti-53BP1(NB100-904, Novus Biologicals, 1:1000 dilution), anti-PML (sc966, Santa Cruz, 1:50 dilution), anti-POLD3 (H00010714-M01, Abnova, 1:100 dilution). For IF-FISH, coverslips were first stained with primary and secondary antibody, then fixed again in 4% formaldehyde for 10 min at room temperature. Coverslips were then dehydrated in an ethanol series (70%, 80%, 90%, 2 minutes each) and incubated with 488-telG PNA probe (Panagene) at 75 °C for 5 min and then overnight in a humidified chamber at room temperature. Coverslips were then washed and mounted for imaging.

### Image acquisition

For live imaging, cells were seeded on 22×22mm glass coverslips (no. 1.5; Fisher Scientific) coated with poly-D-lysine (Sigma-Aldrich) in single wells of a 6-well plate. When ready for imaging, coverslips were mounted in magnetic chambers (Chamlide CM-S22-1, LCI) with cells maintained in L-15 medium without phenol red (Invitrogen) supplemented with 10% FBS and 1% penicillin/streptomycin at 37 °C on a heated stage in an environmental chamber (Incubator BL; PeCon GmbH). Images were acquired with a spinning disk confocal microscope (DM4000; Leica) with a 100x 1.4 NA objective, an XY Piezo-Z stage (Applied Scientific Instrumentation), a spinning disk (Yokogawa), an electron multiplier charge-coupled device camera (ImageEM; Hamamatsu Photonics), and a laser merge module equipped with 488 and 593 nm lasers (LMM5; Spectral Applied Research) controlled by MetaMorph software (Molecular Devices). Images were taken with 0.5 μm spacing for a total of 6 μm and 5 mins time interval for 2-4 hours for both GFP and mCherry channels. Fixed cells were imaged using a 100x 1.4 NA objective on an inverted fluorescence microscope (DM6000, Leica Micro- systems) equipped with an automated XYZ stage (Ludl Electronic Products), a charge-coupled device camera (QuantEM 512SC, Photometrics), an X-LIGHT Confocal Imager (Crisel Electrooptical Systems) and an IDI high performance fluorescence illuminator equipped with 405, 445, 470, 520, 528, 555 and 640 nm lasers (89 North and Cairn Research LTD), controlled by Metamorph Software (MDS Analytical Technologies). Images were taken with 0.3 μm spacing for a total of 8 μm.

### Image processing

All images shown are maximum-intensity projections from all slices in z-stacks generated in Image J (Schneider et al., 2012). Quantifications of images and plotting of figures were done in MATLAB (MathWorks). For live imaging, TRF1 foci in the GFP channel were identified with a 3D bandpass filter with custom MATLAB code modified based on gift code from Stephanie Weber (Berry et al., 2015). The number of segmented TRF1 foci and integrated fluorescence intensity per foci were calculated at each time point. The integrated fluorescence intensity per foci was calculated by first summing up the total intensity over all Z slices in the foci and then calculating the average value over all foci in the cell. For colocalization analysis of fixed images, both channels were segmented with a 3D bandpass filter. The number of colocalized foci and the total fluorescence intensity summed over all Z slices and over all colocalized foci in one cell were plotted.

### Statistical analyses

All p values were generated with two-sample t-test in MATLAB with function ttest2.

## Competing interests

non.

